# UPR deficiency in budding yeast reveals a trade-off between ER folding capacity and maintenance of euploidy

**DOI:** 10.1101/2024.11.22.624941

**Authors:** Constantine Bartolutti, Allison J Kim, Gloria A Brar

## Abstract

The Unfolded Protein Response (UPR) was discovered in budding yeast as a mechanism that allows cells to adapt to ER stress. While the Ire1 branch of this pathway is highly conserved, it is not thought to be important for cellular homeostasis in the absence of stress. Surprisingly, we found that removal of UPR activity led to pervasive aneuploidy in budding yeast cells, suggesting selective pressure resulting from UPR-deficiency. Aneuploid UPR-deficient cells grew better than euploid cells, but exhibited heightened general proteostatic stress, a hallmark of aneuploidy in wild-type cells. Modulation of key genes involved in ER proteostasis that were encoded on aneuploid chromosomes, could phenocopy the effects of aneuploidy, indicating that the reason cells require UPR activity to maintain euploidy is to counteract protein folding stress in the ER. In support of this model, aneuploidy in UPR-deficient cells can be prevented by expression of a UPR-independent general ER chaperone. Overall, our results indicate an unexpected role for the UPR in basal cell growth that is sufficiently important for cells to accept the costly trade-off of aneuploidy in the absence of UPR activity.

## Introduction

The unfolded protein response (UPR) is a conserved cellular pathway that cells activate in response to endoplasmic reticulum (ER) stress, helping them adapt to and cope with the stress. In metazoans there are three main branches of the UPR, one of which is conserved in budding yeast. The ER transmembrane protein Ire1 can be activated by several cues, including those that result from a buildup of misfolded proteins in the ER, subsequently driving splicing of the cytoplasmically retained intron in the mRNA encoding Hac1 (known as XBP-1 in metazoans) (Sidrauski & Walter, 1997). Removal of the intron allows translation of the resultant spliced mRNA (termed *sHAC1*) (Chapman & Walter, 1997). Hac1 protein acts as a transcription factor to activate transcription of genes, including those encoding ER-localized chaperones (such as Kar2, known as BiP or GRP78 in mammals) (Cox and Walter, 1996; Mori et al., 1992; Mori et al., 1996). Targets like Kar2 increase the capacity of the ER to fold proteins and allow the cell to inactivate the UPR and return to homeostasis.

Most studies of UPR mechanism have relied on harsh external triggers, including drugs such as dithreothreitol (DTT) or tunicamycin. The UPR is also known to be important under physiological conditions in metazoans, including during development and B-cell differentiation (Reimold et al., 2000; Reimold et al., 2001). However, the UPR is not thought to function in maintaining basal homeostasis in yeast or human cells. Consistently, UPR deficiency has not been reported to result in aneuploidy. By adaptive aneuploidy, fitness of slow-growing cells is improved by gain or loss of specific chromosomes leading to population-wide loss of euploidy. There is increasing understanding that this can occur efficiently in response to specific genetic or environmental conditions in budding yeast, an organism that is typically studied in its stable euploid state (Chen et al., 2012).

Organisms rely on the proper balance of gene expression for optimal fitness, a balance that is maintained by stable euploid chromosome content. Aneuploidy results in a widespread imbalance in gene expression and has been shown to generally lead to poor cellular fitness compared to euploidy (Torres et al., 2007; Niwa et al., 2006; Hughes et al., 2000; Williams et al., 2008). This is thought to occur, at least in part, because protein complexes depend on precise stoichiometry of their protein components. Because transcription and translation generally occur in proportion to genomic copy number in aneuploid cells, degradation of over-synthesized protein complex components is required to prevent aggregation of orphan subunits (Torres et al., 2007; Pavelka et al., 2010; Springer et al., 2010; Tang et al., 2011). These excess subunits tax the cellular machinery that promotes protein folding and degradation in aneuploid cells (Oromendia et al., 2012). Although overtaxed proteostasis machinery leads to poor growth of aneuploid cells as a rule, there are several notable exceptions to this general principle. Specific stress conditions select for aneuploid cells from the population if they increase the expression of key genes needed for mitigation of stress (Pavelka et al., 2010; Chen et al., 2012). However, adaptive aneuploidy would not be expected during disruption of a pathway without important basal homeostatic function.

In this study we unexpectedly identified consistent and robust emergence of specific aneuploidies in UPR-deficient yeast cells. The aneuploidies that we observed in UPR-deficient haploid and diploid yeast cells were, remarkably, beneficial for growth, even in the absence of ER stressors. This benefit of aneuploidy occurs despite the presence of general proteostatic stress that is typical of aneuploidy in otherwise UPR-proficient cells. We find that maintenance of euploidy under mitotic conditions requires continuous presence of the UPR, which we propose reflects a previously unrecognized requirement for basal UPR activity for cell fitness. Genes that lead to improved ER proteostasis seem to drive these aneuploidies, which can be prevented by increased levels of ER chaperones. Our findings indicate that baseline ER functions in protein folding require UPR activity, and that loss of this activity is sufficiently costly to cells that they adapt by becoming aneuploid, at the expense of general proteostasis.

## Results

### UPR-deficient budding yeast cells become aneuploid in the absence of external ER stress

To facilitate studying the UPR, we created multiple diploid SK1 budding yeast strains homozygously deleted for the gene encoding the conserved UPR sensor Ire1. To our surprise, although wild-type (WT) strains are genomically stable, whole genome sequencing (WGS) of populations of *ire1Δ* cells revealed aneuploidy in all cases. Three independently generated *ire1Δ* diploid strains lacked one copy of chromosome V and one strain gained an extra copy of chromosome X (Figure 1A, 1B). Aneuploid cells dominated these populations, with 0.55X as many chromosome V WGS reads in the UPR-deficient cells compared to WT, for example (Figure 1A). We asked whether this was generally related to UPR function by performing WGS of populations of two independently generated *hac1Δ* diploid strains. WGS confirmed that all *hac1Δ* diploid strains also were aneuploid. For one, we again observed loss of one copy of chromosome V (Figure 1A, 1B); the other lost chromosome VIII and gained chromosome II (Figure 1A, 1B).

**Figure 1.**
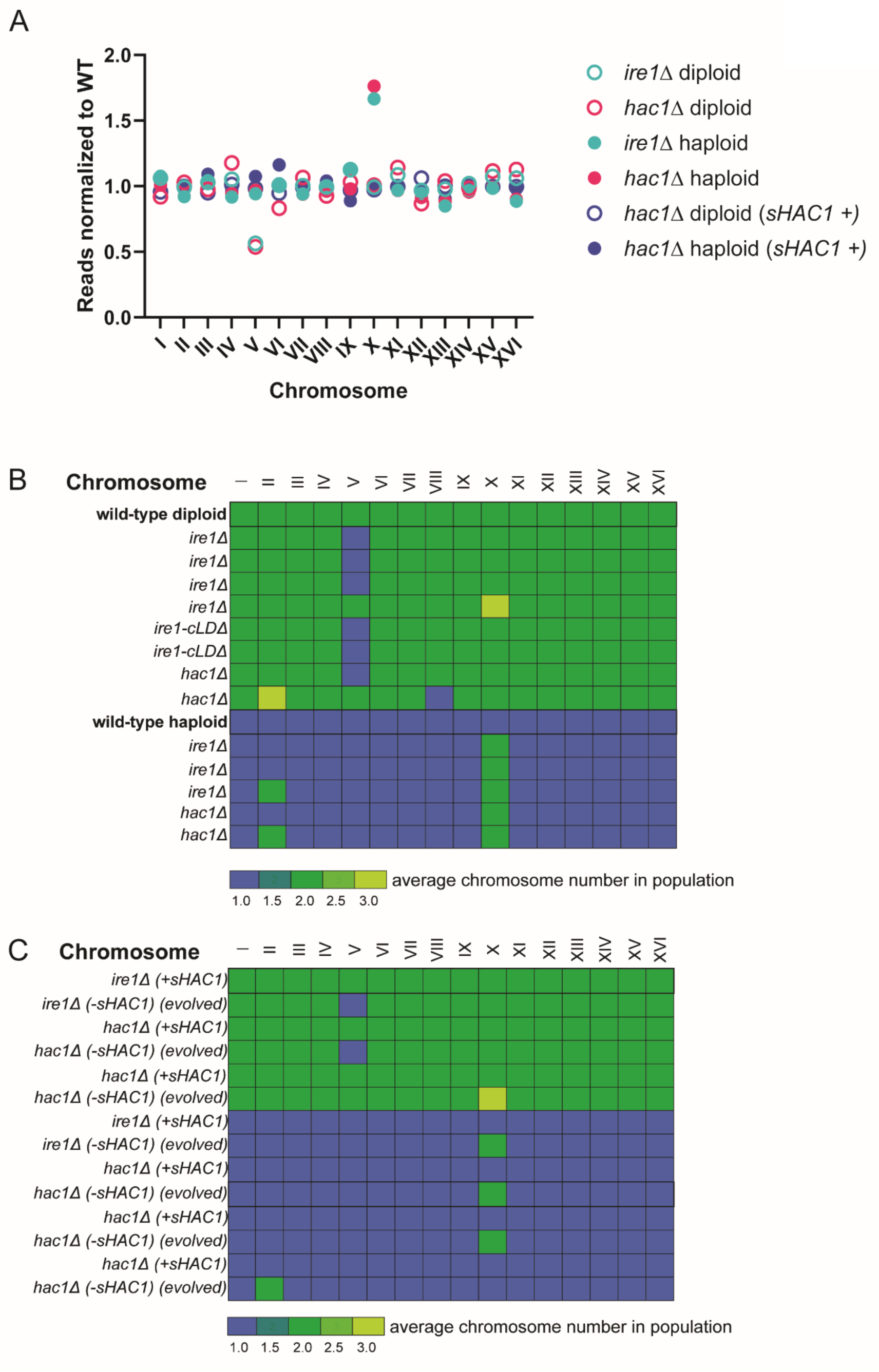
UPR deficient cells robustly become aneuploid for specific aneuploidies. (A) WGS data plotted by chromosome on x-axis for UPR-deficient strains with or without Hac1 expression. Percentage of total aligned WGS reads is plotted relative to a known wild type euploid. Data for representative strains is shown. *sHAC1* + strains carry aTC-driven Hac1 and are grown in the presence of aTC (B) A chart of all WGS-characterized UPR-deficient strains. (C) A table documenting all WGS-characterized evolved strains. Evolved strains are directly below the strain they were evolved from. (B-C) Ploidy for each chromosome is rounded to the nearest half-integer.

Several studies examining budding yeast cells deleted for either *HAC1* or *IRE1* have been performed, and aneuploidy associated with these mutants has not previously been reported; however, all studies known to us previously examined haploid yeast strains. To determine whether aneuploidy in UPR deficient strains was diploid-specific we performed WGS on haploid *ire1Δ* and *hac1Δ* strains. We found no diploid-specific aneuploidy, with three independent *ire1Δ* haploid strains and three independent *hac1Δ* haploid strains displaying pervasive aneuploidy (Figure 1A, 1B). These aneuploidies invariably involved gain of chromosome X and in two cases, we also observed gain of chromosome II (Figure 1B). As with diploids, these aneuploid cells were dominant in the population, as gauged by the approximately 1.8X WGS reads observed for chromosome X in haploid *ire1Δ* or hac*1Δ* cell populations, relative to WT controls (Figure 1A).

Because haploid cells are inviable if any chromosome is lacking, it was not surprising that loss of chromosome V was not observed. We noted that all UPR-deficient strains that we analyzed via WGS were revealed to be aneuploid, regardless of strain generation technique (see methods).

### Hac1 expression prevents aneuploidy driven by UPR deficiency

To confirm that the aneuploidy that we consistently observed in cells deleted for UPR genes was due to loss of UPR activity, we constructed new *ire1Δ* and *hac1Δ* strains in a similar manner to our previous effort, but this time we also suppled cells with s*HAC1* mRNA under control of a tetracycline-inducible promoter (Azizoglu et al., 2021). Following transformation with selection cassette replacement constructs targeted to the *HAC1* or *IRE1* genomic locus, cells were grown on agar plates containing anhydrotetracycline (aTC). In total, we constructed three independent strains for each ploidy and genotype (*ire1Δ* and *hac1Δ*, and haploid and diploid) by this strategy. As expected, we observed that hac*1Δ* strains supplied with Hac1 through this strategy were no longer prone to aneuploidy, as assessed by analysis of three haploid and three diploid strains, all of which resulted in WGS data consistent with euploid chromosome content (example in Figure 1A). Moreover, Hac1 supplementation prevented aneuploidy in all three haploid and all three diploid strains we analyzed that were deleted for *ire1Δ* (example in Figure 1A), indicating that loss of Hac1 activity was the cause of the widespread aneuploidy in *ire1Δ* and *hac1Δ* cell lines.

The ER UPR is not thought to be functionally important in mitotically dividing cells in the absence of external triggers of ER protein folding stress, like DTT or tunicamycin. Thus, we wondered if the aneuploidy was a result of the stress associated with DNA transformation used to generate *ire1Δ* and *hac1Δ* strains, which requires permeabilization of cells and homology-directed repair. To determine whether this was the case, we tested whether euploid UPR-deficient strains, generated in the presence of aTC-induced Hac1 expression, would become aneuploid when grown in the absence of aTC. Indeed, for 7 independent *ire1Δ* or *hac1Δ* strains that were streaked to single colonies on agar plates lacking aTC (including both haploids and diploids for either genotype), we observed emergence of aneuploidy in all cases (Figure 1C). Moreover, the patterns of aneuploidy observed were similar to those identified in *ire1Δ* and *hac1Δ* transformants, with 4 instances of gain of chromosome X, 2 instances of loss of chromosome V, and 1 instance of gain of chromosome II (compare Figure 1A and Figure 1C). We concluded that UPR activity is crucial to maintain euploidy, even in the absence of conditions in which UPR activation is thought to occur.

### Aneuploid UPR-deficient cells exhibit better growth than euploid UPR-deficient cells

The reproducible loss of euploidy in UPR-deficient cells suggested a selective advantage for chromosome V loss in diploid cells and chromosome X gain in haploid cells lacking UPR activity. We measured doubling times for either euploid or aneuploid diploid or haploid *hac1Δ* or *ire1Δ* cells in two rich growth media types: the glucose-containing YPD (Figure 2B) or BYTA (Figure 2C), which lacks a fermentable carbon source and instead contains acetate. We observed reproducibly faster doubling times in the aneuploid UPR-deficient cells, with ∼3x shorter doubling times for haploids and ∼2x faster growth for diploids, compared to UPR-deficient euploid cells (Figure 2A, B). Supporting the idea that a lack of UPR activity drives the growth advantage observed in aneuploid cells, we observed an even more pronounced difference in the presence of externally induced UPR stress. UPR-deficient aneuploid cells displayed a ∼5X shorter doubling-time, relative to UPR-deficient euploid cells in YPD media containing 1mM DTT (Figure 2D).

**Figure 2.**
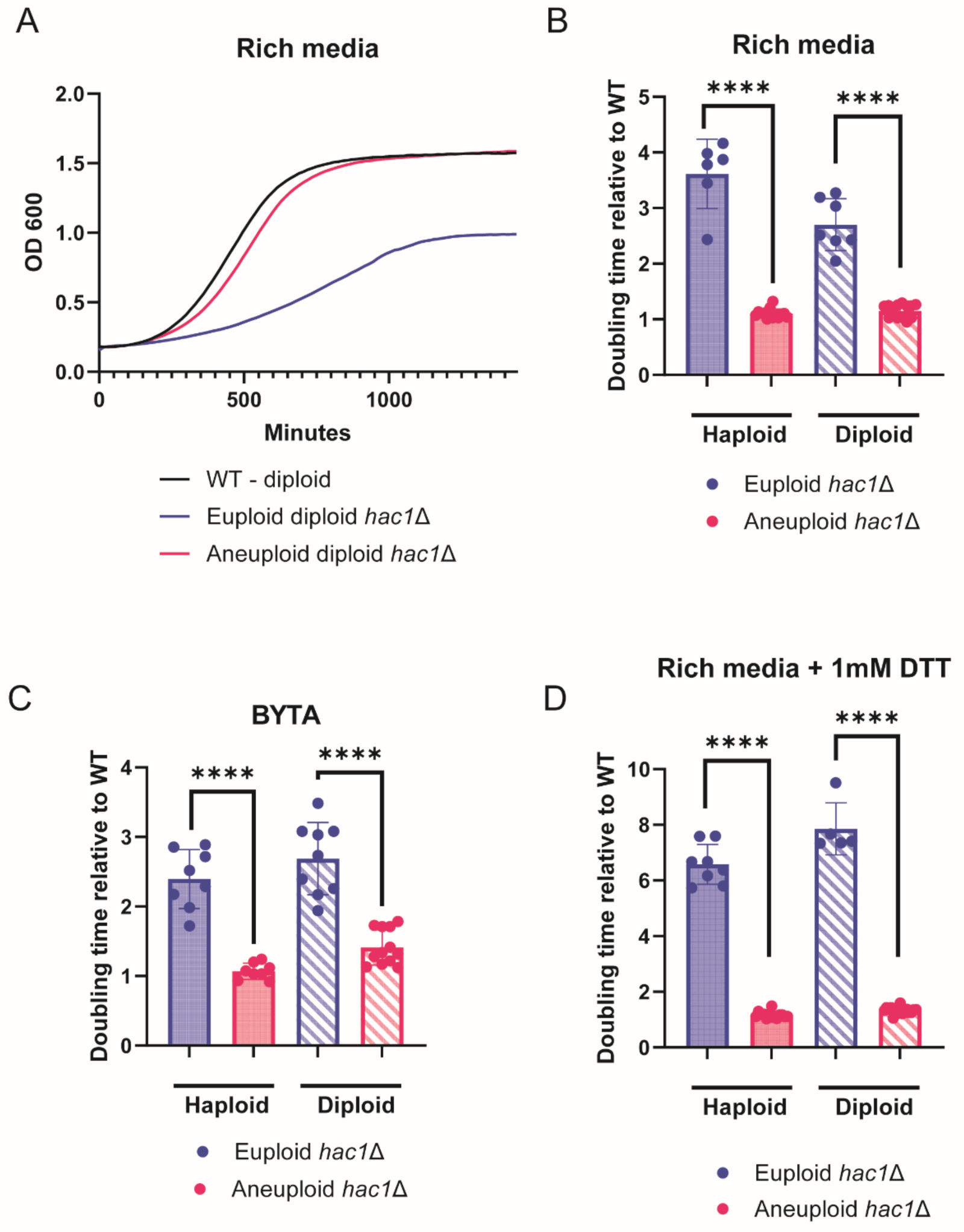
UPR-deficient aneuploidies provide a growth advantage over euploid strains. (A) Representative growth curve shown for WT and UPR-deficient strains. (B-D) Doubling times generated from growth curves as in (A). Strains grown in rich media (YPD) (B), nonfermentable carbon source media (BYTA) (C), and rich media with ER stressor (YPD + 1mM DTT) (D). Doubling times are normalized to wild type for each condition. Each datapoint represents averaged replicates. The haploid aneuploid strain in each case is a representative chromosome X gain strain, while the diploid aneuploid in each case is a representative chromosome V loss strain. Minimum of 5 biological replicates per experiment. Significance is determined from unpaired t-test with Welch’s correction. P-value significance is **** < .0001.

### Aneuploid UPR-deficient cells exhibit the characteristic proteostatic cost of chromosomal imbalance

Prior studies focusing on aneuploidy have reported a cellular cost to chromosome imbalances that results, at least in part, from increased flux through the ubiquitin proteasome system due to the need to degrade the excess subunits of protein complexes that are synthesized in improper stoichiometry (Torres et al., 2007; Pavelka et al., 2010; Springer et al., 2010; Tang et al., 2011) (Oromendia et al., 2012). Consistent with prior work in haploid cells containing one additional chromosome (Torres et al., 2007), both diploid *ire1Δ* and *hac1Δ* cells lacking one copy of chromosome V express transcripts in proportion to chromosome copy number. Specifically, mRNA-seq analysis revealed that mRNAs expressed from chromosome V were present in aneuploid strains at roughly half of the abundance seen in WT strains (Figure 3A, B). To test whether these imbalances in transcript abundance lead to the type of proteostasis defects reported previously in otherwise WT aneuploid strains (Oromendia et al., 2012), we genomically tagged Hsp104 with GFP in either euploid or aneuploid *hac1Δ* cells. Hsp104 is normally diffusely localized throughout the cell, however it also localizes to protein aggregates, if present, making Hsp104 foci a hallmark of proteostatic stress (Liu et al., 2010; Oromendia et al., 2012). We observe a robust increase in cells with prominent Hsp104 foci in the aneuploid populations (Figure 3C-E). Similar results were obtained whether cells were growing in exponential or post-diauxic phase, and for both UPR-deficient haploids (+chromosome X) and diploids (-chromosome V). In sum, these results indicate that, although the aneuploidies resulting from UPR-deficiency improve growth rate relative to euploid UPR-deficient cells, this comes at the cost of increased general proteostatic stress.

**Figure 3.**
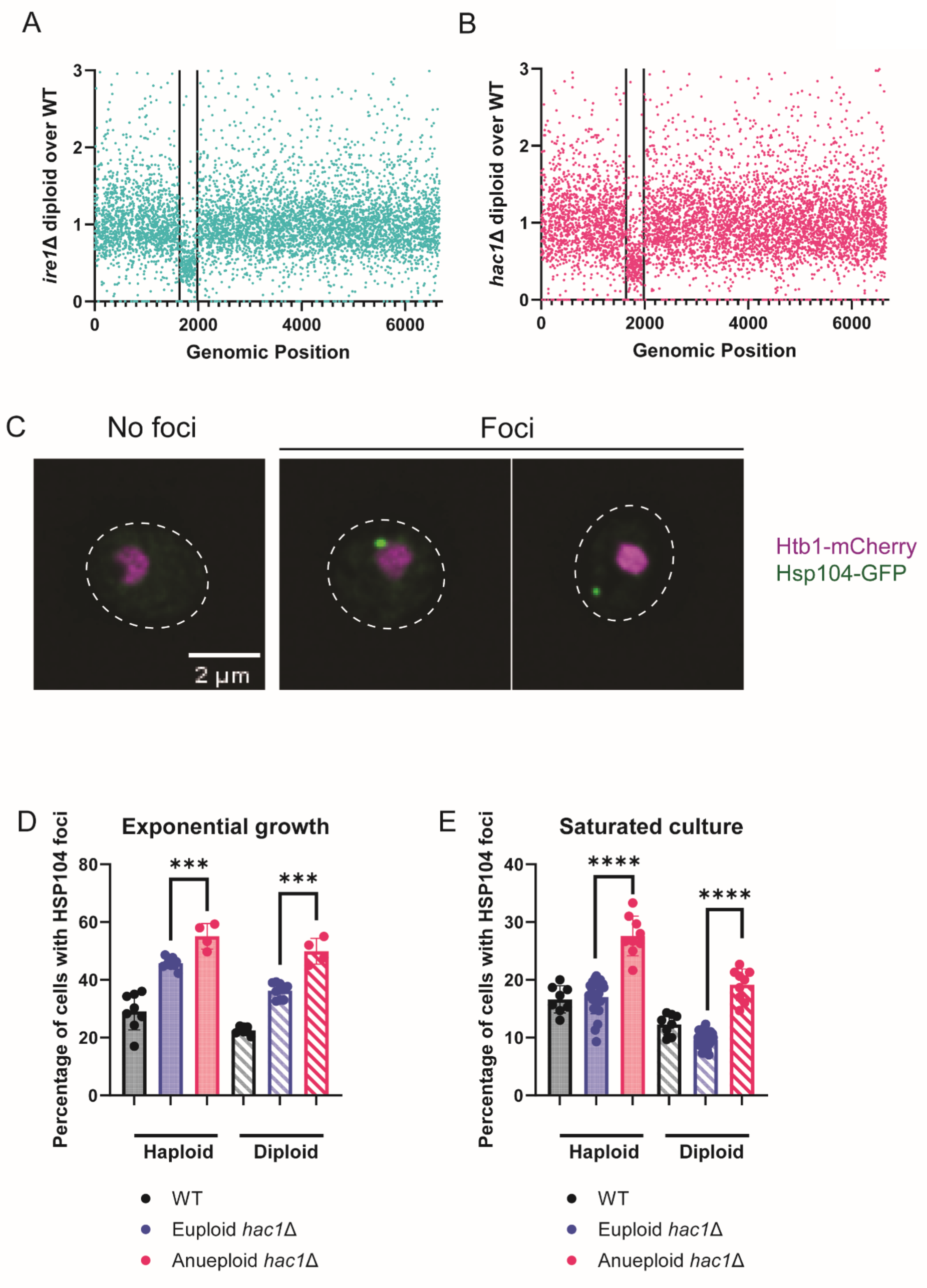
Aneuploid UPR-deficient strains have characteristic proteostasis defects associated with aneuploids. (A-B) mRNAseq reads plotted per gene normalized to wild type for *ire1Δ* (A) and *hac1Δ* (B) deletes. X-axis spans genomic positions by chromosome from I to XVI. Vertical black lines delineate the start and end of chromosome V. (C) Representative single Z slice images for cells with and without Hsp104 foci all taken from aneuploid cells. Two examples of commonly seen foci are given. (D-E) Quantification of cells that have at least one observable Hsp104 foci, regardless of cellular location. Each datapoint represents 300 cells per biological replicate. Cells were counted live while either in exponential growth phase (D) or stationary growth phase (E) in rich media. Minimum of 4 biological replicates for each experiment. Significance is determined from unpaired t-test with Welch’s correction. P-value significance is **** < 0.0001, *** < 0.001.

### Specific ER genes underlie the beneficial impact of aneuploidy in UPR-deficient cells

We observed that UPR-deficiency led to specific patterns of chromosome loss and gain, with loss of chromosome V in diploid cells and gain of chromosome X in haploid cells as the most common pattern (Figure 1B, 1C). Chromosome X contains *KAR2*, the gene encoding the conserved Hsp70 ER chaperone (BiP/GRP78 in humans), which is the best-studied UPR target gene (Figure 4A). We thus hypothesized that expression of extra Kar2 in UPR-deficient cells that have gained a copy of chromosome X may compensate for lack of basal UPR activity and lead to increased growth relative to euploid UPR null cells. To test this, we genomically inserted a single additional copy of *KAR2* under its endogenous regulatory regions to a haploid strain deleted for *HAC1.* This modification alone reduced the doubling time of *hac1Δ* cells in YPD with and without DTT by roughly 3-fold and 5-fold respectively (Figure 4B, C). Remarkably, the doubling time of UPR deficient cells with one extra copy of *KAR2* is comparable to that of haploid cells with an entire extra copy of chromosome X (compare Figures 4B, 4C with 2A, 2C), suggesting that higher expression of Kar2 alone confers the selective advantage offered by this aneuploidy to UPR-deficient cells.

**Figure 4.**
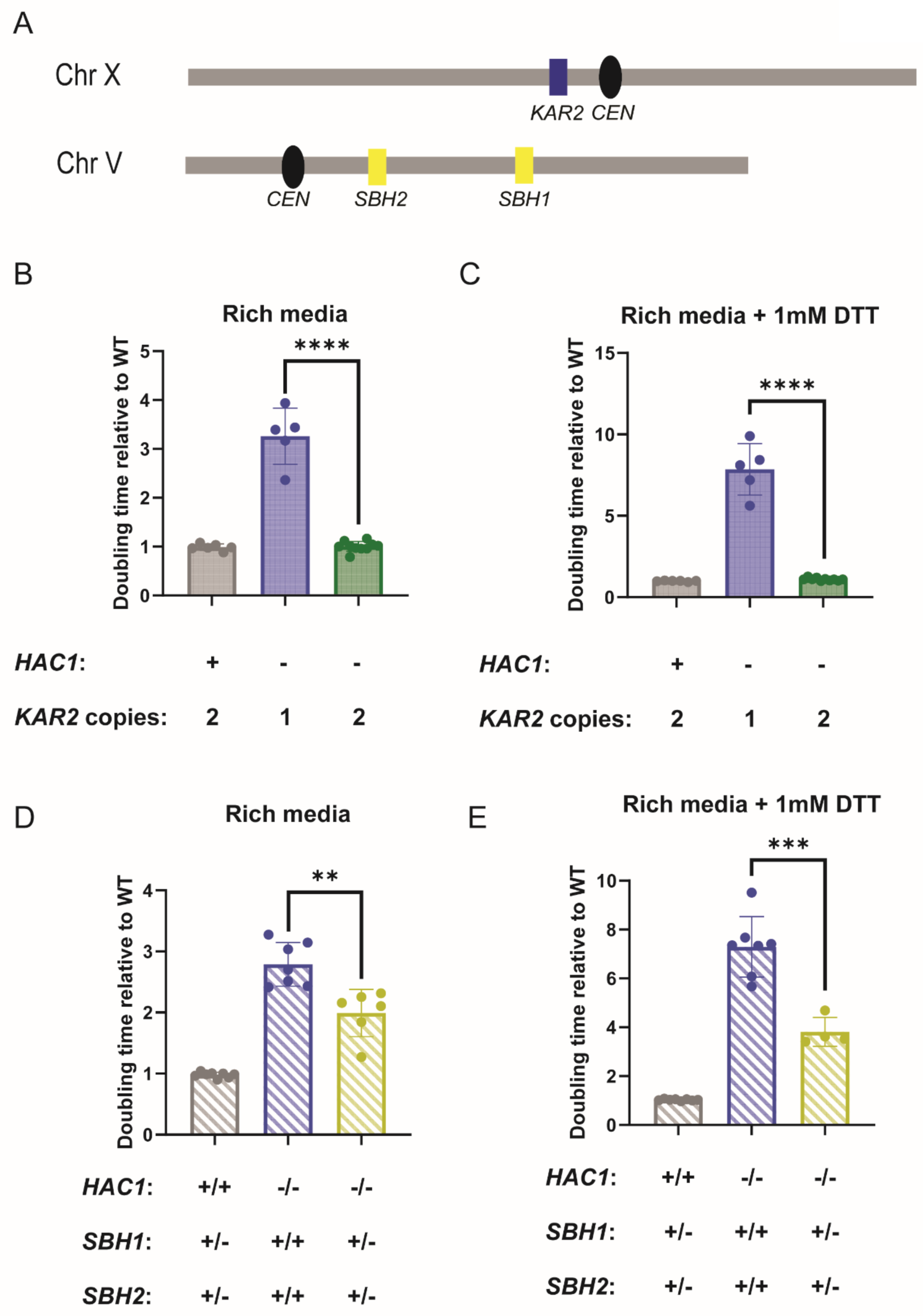
Specific genes underlie the benefit of aneuploid state for UPR-deficient cells. (A) Cartoon representing genomic location of genes tested for UPR-deficient growth rescue. (B-C) Doubling times of haploids were plotted relative to wild type as in Figure 2. Doubling times were plotted for rich media (YPD) (B) and rich media with ER stressor (YPD + 1mM DTT) (C). (D-E) Doubling times of diploids were plotted relative to wild type as in Figure 2. Doubling times were plotted for rich media (YPD) (D) and rich media with ER stressor (YPD + 1mM DTT) (E). Minimum of 4 biological replicates in each experiment. Significance is determined from unpaired t-test with Welch’s correction. P-value significance is **** < 0.0001, *** < 0.001, ** < 0.01.

If Kar2 alone can substitute for UPR activity in supporting normal cell growth, it should be capable of preventing aneuploidy in UPR-deficient cells. To test whether this was the case, we mirrored the strategy we previously used to create euploid UPR-deficient cells (Fig. 1A), but instead of providing cells with aTC-inducible *sHAC1* prior to deletion of *HAC1* or *IRE1,* we generated diploid *ire1Δ* and *hac1Δ* cells in a background expressing aTC-inducible *KAR2* and grew cells on agar media containing aTC. We found that in both *ire1Δ* and *hac1Δ* cells, whether haploid or diploid, this strategy prevented aneuploidy, as determined by WGS analysis of six independent cell populations resulting from independent UPR-deficient strains (Figure 5A).

**Figure 5.**
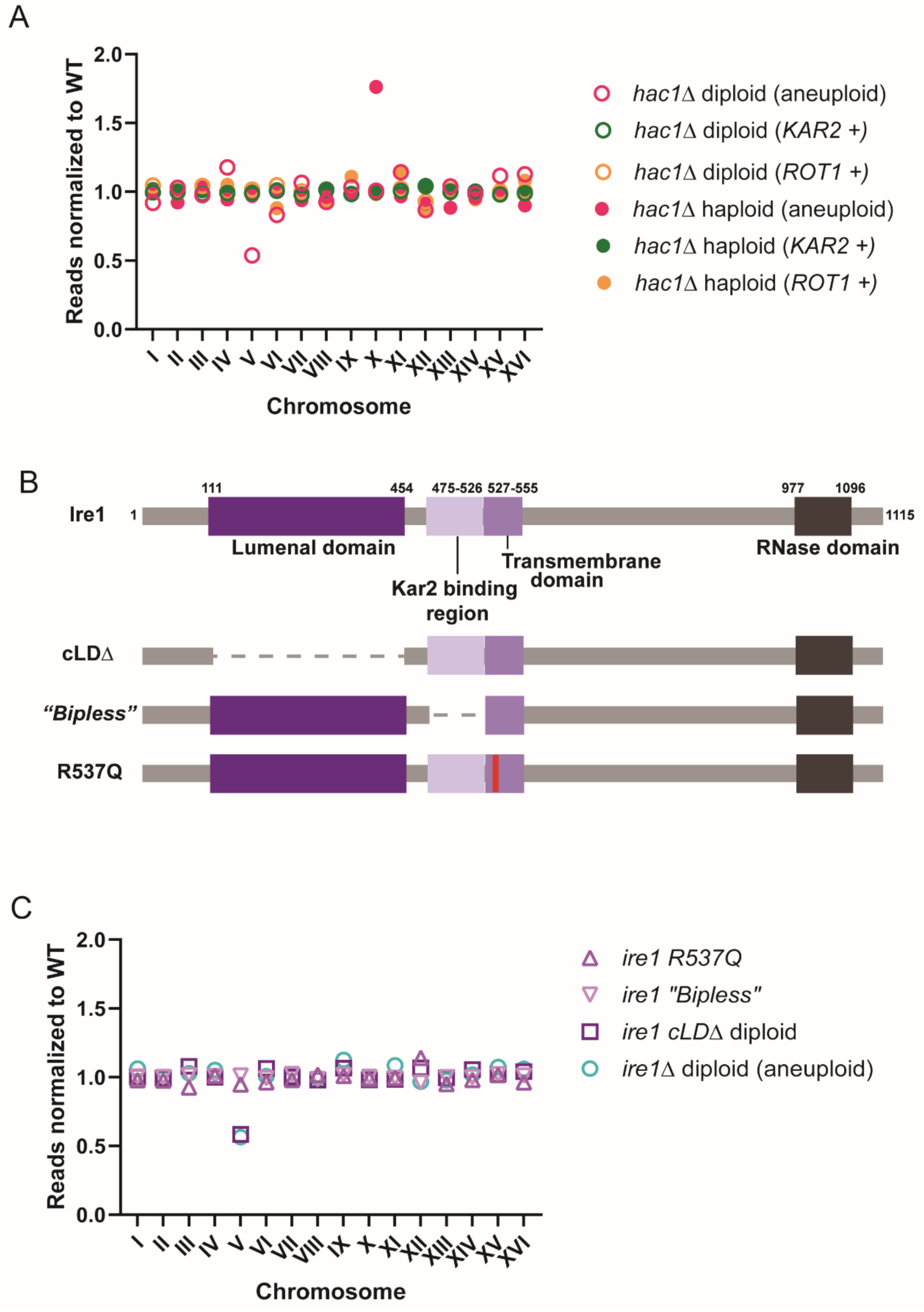
Protein folding defect underlies UPR-deficient strain aneuploidy. (A) WGS data plotted as in Figure 1a. Aneuploid data are the same representative data that was used in Figure 1A, replotted for comparison. *ROT1* + and *KAR2* + strains carried genomically integrated constructs for aTC-driven Rot1 and Kar2, respectively. The *ROT1* + and *KAR2* + data shown are representative data for a single strain for each diploid and haploid. (B) A cartoon depicting relevant domains of Ire1. Representations of mutations made in *IRE1*. (C) WGS data plotted as in Figure 1A. Aneuploid *ire1*Δ diploid is the same representative data from Figure 1a, replotted for comparison. Representative data generated from strains made with the mutations depicted in (B).

Chromosome V contains both *SBH1* and *SBH2*, which encode components of ER translocons through which secretory pathway proteins are inserted into the ER cotranslationally (Figure 4A). We hypothesized that by expressing less of these two proteins, cells could lower flux of proteins into the ER. This would improve fitness under UPR-deficient conditions if overtaxed ER folding is the reason for defective growth in UPR-deficient cells. Consistently, diploid *hac1Δ* cells lacking a single copy of both *SBH1* and *SBH2* displayed a reduced doubling time relative to those with two copies of each (Figure 4C). This was the case in YPD with or without addition of DTT (Figure 4C, 4D). Importantly, the degree of growth benefit was lower than that conferred by the loss of chromosome V, indicating that there are factors beyond these two genes that explain the selective advantage to this particular aneuploidy in UPR-deficient conditions. Nonetheless, based on these experiments, we concluded that adaptive aneuploidies resulting from loss of UPR activity in the absence of exogenous stress are likely to improve growth by improving ER folding capacity, at the expense of general proteostasis.

### ER protein folding stress drives aneuploidy in UPR-deficient strains

Ire1, the UPR sensor, has several well-characterized domains that contribute to its function (Figure 5B). Within the lumen of the ER, there is a 350 amino acid region at the N-terminus of Ire1 that has been shown to directly bind peptides, with a predicted preference for unfolded regions (termed “cLD” here) (Credle et al., 2005; Gardner & Walter, 2011). It is thought that direct binding of unfolded peptides to the cLD results in UPR activation (Credle et al., 2005; Gardner & Walter, 2011). A separate region, between the cLD and Ire1’s transmembrane domain, mediates binding to Kar2/BiP, which is thought to keep Ire1 inactive (Kimata et al., 2003; Pincus et al., 2010). When misfolded ER proteins are present, Kar2 disassociates from Ire1 to bind misfolded regions, which leads to activation of Ire1 (Kimata et al., 2003; Pincus et al., 2010). An *IRE1* mutation that causes loss of the Kar2 interaction is referred to as “*bipless”* and mimics a constitutively active form of Ire1 as Kar2 cannot inactivate Ire1 (Pincus et al., 2010). Ire1’s transmembrane domain has been shown to sense ER membrane integrity, with the mutation of a single amino acid (R537) ablating this activity (Promlek et al., 2011; Halblieb et al., 2017; Ho et al., 2021). Our experiments suggested that the basal role of Ire1 that contributes to aneuploidy prevention is related to reducing misfolded proteins within the ER. We thus predicted that mutating the cLD of *IRE1* would phenocopy deletion of *IRE1*, leading to aneuploidy, whereas loss of membrane integrity sensing or Kar2 binding would not. Using Cas9-mediated mutagenesis we edited *IRE1* at the endogenous locus to either lack the cLD, its Kar2-binding region, or to carry a R537Q mutation. Whole genome sequencing revealed that two independent diploid *ire1-cLDΔ* strains indeed lost chromosome V, whereas *ire1-*R537Q or bipless strains maintained euploidy (Figure 5C). This result suggests that Ire1’s function in peptide binding within the ER is the key UPR role that must be maintained for euploidy.

Considering the importance of the Ire1 region that binds misfolded proteins and the fact that overexpressing *KAR2* or reducing translocon component expression in euploid *hac1Δ* cells improves growth rate, we wondered whether increasing ER folding capacity by means independent of the UPR could compensate for a lack of UPR activity in maintaining euploidy. Toward this end, we supplied cells with aTC-inducible Rot1, an ER chaperone that is not induced by UPR activity. Intriguingly, in the presence of aTC-induced Rot1 we were able to delete *HAC1*, in haploids and diploids, while maintaining euploidy in these populations. This was determined by WGS analysis of 2 cell populations resulting from independent UPR-deficient strains (Figure 5A). We concluded that the UPR plays an important role in supporting ER protein folding, even in basal mitotic growth conditions and in the absence of external sources of ER stress. This role is critical enough for cell fitness that when absent, cells become aneuploid. In doing so, UPR-deficient aneuploid cells synthesize many factors out of proper stoichiometry and overburden their general proteostasis machinery, while counteracting protein misfolding within the ER (Figure 6).

**Figure 6.**
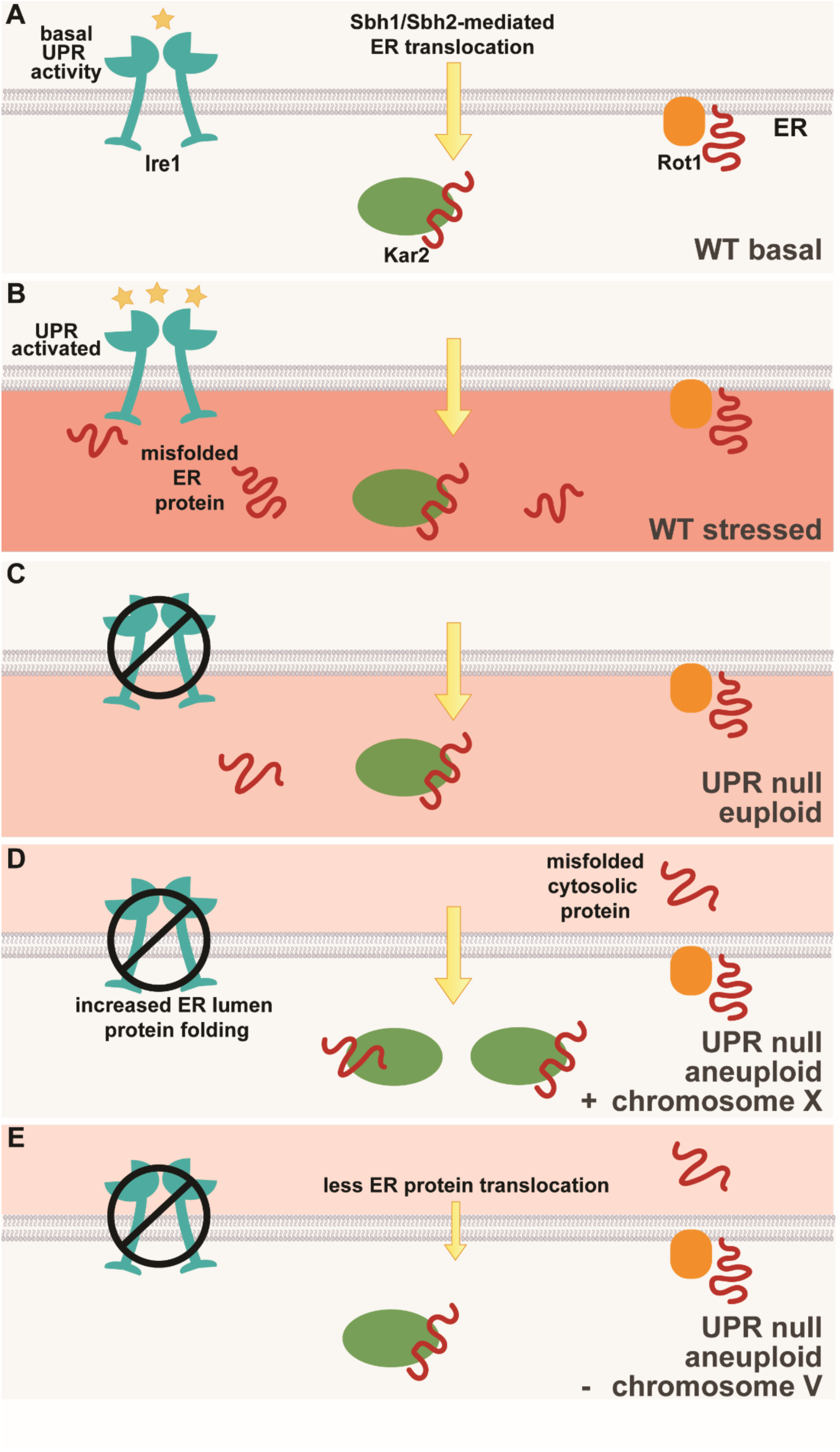
Model for UPR-deficient aneuploidy-based adaptation. (A) In wild-type cells under basal conditions, there is low ER stress that the cell tolerates by activating low levels of UPR activity. (B) In response to severe ER stress, a wild-type cell will activate the UPR strongly. (C) In euploid UPR-deficient cells, the low levels of ER cannot be mitigated by the UPR, leading to poor cellular fitness (D-E) In aneuploid UPR-deficient cells, the chromosome imbalance enhances ER protein folding either via increased ER chaperone expression (D) or decreased relative ER protein import (E), which comes at the expense of mild cytosolic proteostasis deficiency.

## Discussion

We have found that UPR-deficient budding yeast cell populations reliably become aneuploid. These aneuploidies occur in both diploid and haploid cells and are highly specific and characteristic. Haploid UPR deficient cells typically gain a copy of chromosome X, and diploid UPR deficient cells most often lose one copy of chromosome V. The fact that UPR-deficient cell populations become predominantly aneuploid rapidly–for example in the time it takes to either make stable mutants or grow cell colonies on plates–suggests strong selective pressure for loss of euploidy, even in the absence of stressors known to be associated with ER folding defects. Aneuploidy can be increased by chromosome missegregation and by selection for stable inheritance of an aneuploid chromosome content. Our data indicate that the latter is the case with UPR deficiency but do not address whether the former also occurs.

Consistent with a selective advantage for these specific aneuploidies in UPR-deficient cells, we found that either haploid cells disomic for chromosome X or diploid cells monosomic for chromosome V divide significantly more rapidly than euploid controls lacking the UPR. This is the opposite of the pattern seen for otherwise wild-type haploid cells disomic for chromosome X, which divide more slowly than control euploid cells (Torres et al., 2007; Pavelka et al., 2010). The characteristic slow growth of aneuploid yeast cells has led to the conclusion that aneuploidy comes at a cost to cellular fitness. The mitotic growth defect observed in aneuploid cells by systematic studies in yeast has been attributed, in part, to the general proteostatic cost of synthesizing excess protein complex members that must be degraded (Torres et al., 2007; Pavelka et al., 2010; Springer et al., 2010; Tang et al., 2011; Oromendia et al., 2012). This conclusion is based on a number of observations, including expression of genes proportional to their genomic copy number, the increased dependence of aneuploid cells on proteasome activity for their survival and the increased presence of Hsp104 foci in aneuploid relative to wild-type cells (Torres et al., 2007; Pavelka et al., 2010; Springer et al., 2010; Tang et al., 2011) (Oromendia et al., 2012). We observe evidence of similar costs in aneuploid UPR deficient cells, with transcript abundance proportional to genomic copy number and Hsp104 foci elevated in aneuploid versus wild-type cells. This suggests that there is a trade-off occurring in the absence of UPR activity that favors support for ER protein folding at the expense of general proteostasis. It has been recently found that non-laboratory aneuploid yeast strains exhibit a cost as a result of additional factors, including snoRNAs from the extra chromosomes (Rojas et al., 2024). Such a cost would be expected to apply in the case of the UPR-deficient aneuploid cells, although we did not assay this directly in our study.

Several lines of data suggest that the dependence of non-stressed mitotic cells on UPR activity is based on improving ER protein folding. These include the observation that overexpression of either Kar2 (a UPR-induced ER chaperone) or even Rot1 (a UPR-independent ER chaperone) can prevent aneuploidy in UPR-deficient cells. Additionally, a role for supporting ER homeostasis is suggested by the identities of genes on the common chromosomes that are involved in aneuploidy in UPR-deficient cells. Simply doubling *KAR2* copy number was sufficient to phenocopy the growth benefit offered by an extra copy of chromosome X in UPR-deficient cells. This strongly suggests that the primary reason chromosome X gains are so commonly seen among UPR-deficient aneuploidies is that additional Kar2 increases cellular fitness in this context. For diploid cells, the reason that chromosome V loss is beneficial in cells lacking the UPR is still partly unresolved, but our data suggest that two genes encoding ER translocon components on this chromosome, *SBH1* and *SBH2,* are part of the answer. We hypothesize that less Sbh1 and Sbh2 could improve ER protein folding by reducing translocation into the ER, thus providing less competition for ER chaperones. Consistent with this model, diploid UPR-deficient euploid cells that were heterozygous for both genes divide more rapidly than those homozygous for *SBH1* and *SBH2*. However, the degree of growth rescue was less prominent than that seen in diploid UPR-deficient cells lacking the entire chromosome V, implying that other gene(s) on this chromosome may also be important for this role. One additional aneuploidy that we observed in several cases in UPR-deficient cells was gain of Chromosome II, which was sometimes seen in conjunction with other aneuploidies. The reason for selection of an additional Chromosome II remains unclear, but we note that it includes a gene encoding a well-studied ERAD component: Der1, as well as Rot2, which is synthetically lethal with Rot1 mutations (Bickle et al., 1998). Rot2 has not been well studied, but it is located at the ER membrane facing inward into the lumen, like Rot1, suggesting that it may serve a similar role.

The UPR was discovered in budding yeast and has been studied extensively in this organism over decades, yet no previous studies have reported aneuploidies associated with the loss of *HAC1* or *IRE1*. For cases in which global data in such mutants are available, all are in the reference S288C budding yeast strain or its recent derivative W303, and all do not display evidence of aneuploidy. There are several possible reasons that our more recently domesticated strain background (SK1) may reveal a role for the UPR in maintaining euploidy that has not been seen in S288C/W303. First, growth of SK1 may require greater support for ER folding through the UPR than S288C. However, northern blot analysis examining *HAC1* splicing of SK1 WT cells in rich mitotic growth conditions did not reveal detectable levels of spliced *HAC1* (data not shown), similar to observations originally made in S288C. Furthermore, ribosome profiling revealed translation of *HAC1* that was equivalently low in SK1 and W303 cells. Using ribosome profiling, we measured this to be at 59 and 56 reads per kilobase per million (RPKM), respectively, approximately 2.7% of that measured in DTT-treated cells (Van Dalfsen et al., 2017; Thorburn et al., 2013) This very low level of basal mitotic UPR induction in SK1 is interesting given that our data indicate that cells benefit from UPR activity under such conditions, and yet is consistent with the ability of doubling only one UPR target (Kar2) to fully rescue the UPR-deficient growth rate defect.

A second potential explanation for the lack of aneuploidy driven by UPR deficiency in standard laboratory yeast strains is the higher cost of aneuploidy in these strain backgrounds (Hose et al., 2015; Gasch et al., 2016; Scopel et al., 2021). W303, in particular, has been shown to be unusually aneuploid resistant because of a mutation in the gene encoding the RNA binding protein Ssd1 (Hose et al., 2020). However, mutation of *SSD1* in SK1 did not prevent aneuploidy in UPR deficient strains (data not shown, n=3). We find relative *KAR2* expression between S288C/W303 and SK1 to be the most compelling reason for lack of observed aneuploidy in UPR-deficient S288C/W303 strains. Both mRNA and translation measurements show that *KAR2* levels are twice as high basally in W303 compared to SK1 (Torres et al., 2007; Brar et al., 2012). In essence, our experiments doubling the levels of Kar2 in SK1 make it similar to these highly derived laboratory strains. This suggests that, more generally, eukaryotic cell types that either express less Kar2 or require more due to high secretory protein load would be prone to aneuploidy.

It is unknown whether there is a role for the UPR in maintaining euploidy in mammals. This is an interesting area of future research, given the high incidence of both UPR activation and aneuploidy in cancer (Madden et al., 2019; Pfau & Amon, 2012; Holland & Cleveland, 2012; Targa & Rancati, 2018). Much work has focused on each individually, with a particular interest in the paradox between the growth defects observed in aneuploid yeast strains and the rapid growth of cancer cells, which tend to be aneuploid. It has been proposed that one explanation for this paradox is the cellular context of cancer cells (Pfau & Amon, 2012; Holland & Cleveland, 2012). Our data are interesting with this model in mind, as we observe a switch in the relative fitness of euploid and aneuploid yeast cells depending on whether they have UPR activity or not. Although *IRE-1* and *XBP-1* are not commonly mutated in cancer cells, the high degree of UPR activation in most cancers suggests a higher basal requirement for this pathway than is typical of healthy cells (Madden et al., 2019). Multiple myeloma is the cancer that has been best studied for a reliance on high UPR activity, and the most common associated aneuploidy is trisomy for chromosome 9, the chromosome housing human BiP/GRP78 (Kumar et al., 2012). Whether this aneuploidy is driven by the need for additional ER protein folding support warrants further study in the light of our findings.

## Methods

### Yeast strains and plasmids

All strains used for experiments in this paper are derived from the SK1 strain background and are specified in supplementary table 1. Plasmids used for experiments are listed in supplementary table 2, as well as primers used for strain or plasmid generation in supplementary table 3. The following alleles used for the Hsp104 experiments were first used in previous studies: Htb1-mCherry (Matos et al., 2008), Hsp104-GFP (Ünal et al., 2011). All integration transformations (see below) were done via restriction enzyme cutting of integration plasmids with PmeI.

### Media and growth conditions

All yeast strains were grown in YPD (1% yeast extract, 2% peptone, 2% glucose) for liquid experiments, unless stated otherwise. Strains were grown on YPD plates plus 250ng/ml of aTC for maintaining euploid state. Nonfermentable carbon source media growth experiments used BYTA (1% yeast extract, 2% bacto tryptone, 1% potassium acetate, and 50mM potassium phthalate) as the liquid media.

### Strain generation

Standard PCR based homologous recombination cassette swapping strategies in yeast were used to generate deletions of genes for this study (Longtine et al., 1998). Importantly when attempting to generate deletions of *IRE1* and *HAC1*, both deletions of haploids, as well as heterozygous deletions of diploids were attempted. Diploids were able to be maintained in a euploid state as heterozygotes but not as homozygotes. Attempts to make diploids involved both two sequential rounds of transformation to delete both copies, as well as sporulating heterozygotes and mating spores that carried an episomal plasmid. For the latter protocol, the episomal plasmid was removed following validation of deletions. Cas9-mediated deletion of the lumenal domain of Ire1 was done by co-transforming cells with a selectable Cas9 expression plasmid and a double-stranded repair template.

To generate euploid UPR-deficient strains, deletion of diploids as well as haploids was done via transformation of targeted selection cassettes into strains carrying genomically integrated aTC-inducible alleles of either *HAC1, ROT1,* or *KAR2* integrated. Strains were plated post-transformation on plates with aTC such that all strains were constantly in the presence of UPR activity mimetics. Heterozygous diploids were then dissected to generate haploids to backcross for construction of stable haploid or diploid lines. Importantly, strains were always grown on aTC plates except for when experiments required a no UPR condition.

### DNA extraction for Whole Genome Sequencing

DNA extraction was performed as reported in Morse et al., 2024. Final pellet was resuspended in 100ul DEPC treated water instead of 50ul of TE.

### Whole Genome Sequencing

Extracted DNA was submitted to SeqCenter for Illumina Whole Genome Sequencing. SeqCenter also performed Variant Calling analysis for the samples using the breseq software package. A minimum of 1 million mapped reads were achieved for all samples. For all WGS samples the total number of aligned reads was summed and then a percentage was calculated for number of reads per total aligned reads. Verified euploid strains were then used to normalize the percentages per putative aneuploid strain.

### Growth Curve Analysis

All strains were grown overnight in YPD before being back-diluted to an OD_600_ of .1 in a 96-well plate. Plates were sealed with a Breathe Easy Cover (Sigma-Aldrich), and grown at 30C for 24 hours in a plate reader. Absorbance readings were collected every 15 minutes by absorbance at 600 nm, with agitation before each reading. All experiments were loaded in triplicate in the 96-well plate and individual replicate growth curves were calculated as the average of these triplicates. Doubling times were calculated by calculating the time of growth between an OD_600_ of 0.3 and an OD_600_ of 0.6 to avoid lag phase and stationary phase.

### Fluorescent Microscopy

Cells were either imaged after 2-4 cell cycles following back dilution to OD_600_ 0.1, or imaged from overnight cultures in stationary phase. Live cells were counted for Hsp104 foci visible in any z plane. 300 cells were counted per replicate and the percentage of cells with foci out of 300 was calculated. Images were acquired using a DeltaVision Elite wide-field fluorescence microscope (GEHealthcare, Sunnyvale, CA). Live cell images were generated using a 100x/1.42 NA oil-immersion objective. Images were deconvolved using softWoRx imaging software (GE Life Sciences). GFP was imaged using 100%T, 0.025s EX: 475/28EM: 523/36. mCherry was imaged using 32%T, 0.025s EX:575/25EM: 632/60.

### RNA extraction

Cells were harvested via pelleting liquid culture at 6000 RCF then aspirating media before snap freezing in liquid nitrogen. Cells were thawed on ice and resuspended in 625ul TES buffer (10 mM Tris pH 7.5, 10 mM EDTA, 0.5% SDS). An equal volume of citrate-buffered acid phenol (pH 4.3, P4682, Sigma-Aldrich) was added to cells, and they were incubated at 65C for 1 hour in a Thermomixer C (Eppendorf) shaking at 1400 RPM. After microcentrifugation (20,000 g for 10 min) the aqueous phase was transferred to a second tube with 600ul acid phenol. The aqueous phase was separated by microcentrifugation (20,000 g for 10 min) and transferred to a third tube with 600ul chloroform. The aqueous phase was separated by microcentrifugation (20,000 g for 10 min) and RNA was precipitated in 100% isopropanol with 350 mM sodium acetate (pH 5.2) overnight at –20C. Pellets were washed with 80% ethanol and resuspended in DEPC water.

### mRNA sequencing

Libraries were constructed as reported in Morse et al., 2024. RPKM (Reads per kilobase million) were calculated for all samples.

### Evolution of yeast strains

Verified euploid strains were plated directly on YPD plates containing aTC. After 1 day of growth strains were streaked for single colonies on YPG (Glycerol instead of Dextrose as the carbon source) twice before patching single colonies on YPG. Patches of yeast were frozen and saved in the freezer. From patches, strains were grown as is detailed above overnight before harvesting DNA for WGS.

## Acknowledgments

We thank Nick Ingolia, Audrey Gasch and members of the Brar and Ünal labs for feedback on this manuscript; and Jeremy Thorner, Andy Dillin, Nick Ingolia, and Roberto Zoncu for helpful discussions. This work was supported by NIH funding to GAB (R01AG071869).

Author contributions: C.B. and G.B conceived the study. G.B supervised the study. C.B. and A.K. designed and performed the experiments. C.B. evaluated the data and constructed the figures. and G.B. wrote the original draft of the manuscript and reviewed and edited the manuscript.

## Supplementary Files

**Table 1.** Strains used. All strains used in this study are organized by strain number and genotype. Strains available on request.

**Table 2.** Plasmids used. All plasmids used in this study for strain generation or experiment purposes are organized by name and description. Plasmids are available on request.

**Table 3.** Primers used. All primers used in this study for strain or plasmid generation are organized by description and sequence.

